# Genome-wide protein phylogenies for four African cichlid species

**DOI:** 10.1101/110510

**Authors:** Ajay Ramakrishnan Varadarajan, Rohini Mopuri, J. Todd Streelman, Patrick T. McGrath

## Abstract

**Background:** The thousands of species of closely related cichlid fishes in the great lakes of East Africa are a powerful model for understanding speciation and the genetic basis of trait variation. Recently, the genomes of five species of African cichlids representing five distinct lineages were sequenced and used to predict protein products at a genome-wide level. Here we characterize the evolutionary relationship of each cichlid protein to previously sequenced animal species.

**Results:** We used the Treefam database, a set of preexisting protein phylogenies built using 109 previously sequenced genomes, to identify Treefam families for each protein annotated from four cichlid species: *Metriaclima zebra, Astatotilapia burtoni, Pundamilia nyererei* and *Neolamporologus brichardi.* For each of these Treefam families, we built new protein phylogenies containing each of the cichlid protein hits. Using these new phylogenies we identified the evolutionary relationship of each cichlid protein to its nearest human and zebrafish protein. This data is available either through download or through a webserver we have implemented.

**Conclusion:** These phylogenies will be useful for any cichlid researchers trying to predict biological and protein function for a given cichlid gene, understanding the evolutionary history of a given cichlid gene, identifying recently duplicated cichlid genes, or performing genome-wide analysis in cichlids that relies on using databases generated from other species.

## BACKGROUND

The rapid decrease in sequencing costs and the development of broadly applicable genetic tools like TALENs and CRISPR/Cas9 has facilitated the development of a large number of new species as model organisms (Joung and Sander 2013, Doudna and Charpentier 2014, Hsu, Lander et al. 2014,Nemudryi, Valetdinova et al. 2014). For evolutionary biologists, this has been especially fruitful - species with unique evolutionary traits can now be used as model organisms to identify and understand the underlying genetic and cellular mechanisms responsible for trait changes (Goldstein and King 2016). For example, threespine sticklebacks have long fascinated evolutionary biologists for their coexisting phenotypically divergent forms including freshwater/anadromous pairs (Hagen 1967, Mcphail 1969,McKinnon and Rundle 2002). Freshwater lakes created after the retreat of Pleistocene glaciers have been populated by marine sticklebacks, evolving repeated changes in a number of traits. These adaptations include morphological changes to body shape, pigmentation changes, salt handling, and reproductive related behaviors (Bell and Foster 1994,McKinnon and Rundle 2002). A combination of quantitative genetics and resequencing of individuals isolated from freshwater and saltwater habitats identified a large number of loci putatively responsible for evolution of marine-freshwater ecotypes (Colosimo, Peichel et al. 2004, Chan, Marks et al. 2010,Jones, Grabherr et al. 2012). An important conclusion from this research, and a number of other individual examples (Martin and Orgogozo 2013), is that despite the large number of genes that control a trait, natural selection can act in predictable ways, isolating genetic changes in preferred genes in response to specific environmental shifts. An important goal now is to identify additional examples of repeated evolution, and understand why particular genes are repeatedly selected.

Cichlid fishes offer an attractive avenue for this type of research. Cichlids are well-known for their adaptive radiations in the Great Lakes of East Africa. The three largest radiations in Lakes Victoria, Lake Malawi, and Lake Tanganyika have generated between 250 – 500 species per lake in a period of time that ranges from 100,000 to 12 million years (Kocher 2004,Brawand, Wagner et al. 2014). These radiations resulted in exceptional phenotypic diversity in behavior, neurodevelopment, body shape, sexual traits, and ecological specialization. However, due to the speed of evolution, nucleotide diversity between these species is on the order of nucleotide diversity within the human population (Loh, Bezault et al. 2013,Brawand, Wagner et al. 2014). Further, genetic barriers have not formed in this short period, allowing for genetics - phenotypically-divergent species can still interbreed. These peculiarities of the cichlid family make genomics and quantitative genetics approaches particularly attractive. Genes responsible for phenotypic diversity can be identified using quantitative mapping approaches in progeny of intercrossed species, association mapping in outbred animals, or tissue-specific transcriptomics in behaving animals. To facilitate these approaches, high-quality genomes for five cichlid fishes were generated (Brawand, Wagner et al. 2014). It is anticipated that genetic variants and genes responsible for a variety of interesting trait differences will be identified in the coming years.

Due to the difficulty of experimental study of cichlids in the laboratory, assignment of molecular and biological function to genes relies almost exclusively on homology to proteins characterized biochemically, or in model organisms such as *Caenorhabditis elegans, Drosophila melanogaster, Danio rerio,* or *Mus musculus.* Homologous proteins share a common evolutionary ancestry (Fitch 1970), suggesting shared biochemical and/or biological role, justifying the use of homology to assign function to genes identified in cichlid fish. Proteins with shared homology can be characterized as orthologs (which diverged from a common ancestor due to speciation) or paralogs (which diverged from a common ancestor due to a gene duplication event). In general, orthologs are expected to retain similar (if not identical) function with each other. Paralogs are expected to acquire novel function and/or biological roles. For cichlids, paralogs are thought to be especially relevant to their evolution - the cichlid lineage has undergone an increased rate of gene duplication, suggesting that these novel genes could serve important roles in the cichlid’s adaptive radiations (Lynch and Conery 2000,Brawand, Wagner et al. 2014). Cichlids also belong to the teleost infraclass of fish, whose ancestors have undergone a genome-wide duplication event resulting in the duplication of a large number of genes (Taylor, Braasch et al. 2003). Gene duplication can allow resolution of adaptive conflict by allowing a bifunctional ancestral gene to resolve into two specialized genes (Lynch and Force 2000). These gene duplicates have been proposed to play a role in the evolutionary success of the teleost fish, which make up ~96% of all fish. Phylogenetic relationships could potentially be used to identify the cichlid genes that have undergone subfunctionalization. For all of these reasons, it would be helpful to place each cichlid protein into a phylogeny to aid in predicting the gene function for a given cichlid gene.

In this report, we utilized the TreeFam database of protein phylogenies to create protein phylogenies for all completely sequenced cichlid genomes. We analyzed these phylogenies to determine evolutionary relationships for each of these cichlid genes. This data is available for download or searching on a web server, and should be useful to any researchers studying cichlid fish.

## METHODS

### Overview

We employed a phylogeny-based approach to study the function and evolution of <genes of interest> taken from four East African cichlid species. Our aim was to assign each cichlid gene to a pre-defined gene family to identify homologous proteins and their evolutionary relationship. To accomplish this, we used TreeFam, a database of phylogenetic trees drawn from 109 animal genomes (Li, Coghlan et al. 2006, Ruan, Li et al. 2008,Schreiber, Patricio et al. 2014). A webserver implementing the Treefam pipeline is provided (www.treefam.org) to add new proteins of interest to existing TreeFam trees. We implemented this pipeline locally to perform this on a genomic basis.

### Datasets and TreeFam analysis

Protein coding sequences and annotation files for four cichlid species, *A. burtoni, M. zebra, N. brichardi,* and *P. nyererei,* were obtained from the supplemental dataset from the genome sequencing paper (Brawand, Wagner et al. 2014). An improved genome for *M. zebra* was also recently published; protein coding sequence and genome annotation files from this paper were downloaded from NCBI (Conte and Kocher 2015). Annotation files were parsed using custom Python scripts and used to identify the longest protein isoform and amino acid sequence for each gene. This was done to limit the phylogeny to one representative protein isoform for each gene. To assign each of these proteins to a single TreeFam family, we utilized the í ê É ÉÑ ~ ã ëÅ ~ åKéä script provided as part of the TreeFam API (Schreiber, Patricio et al. 2014). This script uses the program HMMER to identify matches using hidden Markov model profiles generated for each of the TreeFam families (Eddy 1998). After this had run on all of the proteins, we collected all of the protein sequences that best matched a given TreeFam to add these to the preexisting phylogeny. Multiple sequence alignments and phylogenies for each TreeFam were retrieved from a locally cloned SQL database with API utilities provided by TreeFam. We used MAFFT (version 7.221) to add the new cichlid proteins to the retrieved multiple sequence alignment using the - ~ÇÇ, - êÉç êÇÉê, and - ~åóëóãÄçä options (Katoh, Misawa et al. 2002,Katoh and Standley 2013). The aligned output was then used to add the new proteins to the retrieved phylogeny file using RaXML (version 8.1.15) using the GAMMA model for rate heterogeneity with the WAG substitution matrix (Stamatakis, Ludwig et al. 2005,Stamatakis 2006).

### Identification of closest relationships to human and zebrafish proteins

For each cichlid protein, we used custom Python scripts to identify the closest human and zebrafish protein using the phylogenetic tree produced by RAxML. The structures of each tree were analyzed using the ETE toolkit, which provides a Python framework for analysis and visualization of protein trees (Huerta-Cepas, Serra et al. 2016). Trees were rooted using a midpoint outgroup method implemented by the ÖÉí | ãá Çéçáåí | çìí Öêçì é function. To find the closest human protein and its evolutionary relationship with a cichlid protein of interest, the trees were then traversed to identify the smallest subtree containing the cichlid protein and one or more human protein. If such a subtree could not be found (i.e. there was no human protein in the phylogeny), the relationship was defined as kçeçãçäç Ö. If the subtree contained a single human protein and a single cichlid protein from the cichlid species, the relationship was defined as I êí Üçäç Ö. If the subtree contained a single human protein and exactly two cichlid proteins from the cichlid species, this relationship was defined as a e~äÑ êí Üçäç Ö with the human protein. Finally, if the subtree contained multiple human proteins, or more than two cichlid proteins from the cichlid species, this relationship was defined as a m~ê~äç Ö. The closest human protein was identified using the shortest branch length. To convert the Ensembl protein ID’s of the human proteins to HGNC identifiers (Gray, Yates et al. 2015), we downloaded mapping data from Ensembl BioMart (Aken, Ayling et al. 2016). An essentially identical process was also performed between all cichlid proteins with zebrafish proteins. An excel spreadsheet (one per species) was then created for each cichlid gene for this information.

PDFs of the resulting phylogenies were rendered using the ETE toolkit. A full size version of each TreeFam phylogeny was created using all species. In addition, a smaller PDF was created from a pruned tree containing a limited number of well-characterized species (human (*H. sapiens*), mouse (*M. musculus*), zebrafish (*D. rerio*), fruit fly (*D. melanogaster*), and nematode (*C. elegans*)), the closely related Nile tilapia (*O. niloticus*), and the four new cichlid species.

## RESULTS AND DISCUSSION

### Identification of human and zebrafish relationships for each cichlid gene

The cichlids species of East Africa have become a popular genomic model to understand the evolution of a number of traits, including differences in morphology, coloration and behavior. To broaden our understanding of the function and evolutionary history of the genes that are encoded in the genomes of four recently-sequenced cichlid species, we performed phylogenetic analysis using the previously published TreeFam pipeline to add the new cichlid proteins to preexisting protein phylogenies generated from a large number of animal species (**Figure 1**). The most current version of the TreeFam database (Schreiber, Patricio et al. 2014), which contains 15,736 phylogenetic trees generated from 109 animal genomes covering ~2.2 million sequences, can be used to study evolutionary relationships between homologous proteins. While this database already includes the African cichlid *O. niloticus* (Nile tilapia), it does not contain four recently sequenced African cichlids: *M. zebra* from Lake Malawi, *P. nyererei* from Lake Victoria, *N. brichardi* from Lake Tanganyika, and *A. burtoni* found in a variety of African lakes and rivers. For all four cichlid species, the majority of cichlid genes, 82.2% –84.7%, contained a hit to a preexisting TreeFam family (**Figure 2**). Using the resulting phylogenies, we identified the closest human and zebrafish gene along with the evolutionary relationship to the cichlid. These included traditional evolutionary relationships (Ortholog and Paralog) and also a novel evolutionary definition we call HalfOrtholog, to account for the large number of cichlid genes that duplicated in the ancestral teleost lineage and are retained in the extant species.

**Figure 1.**
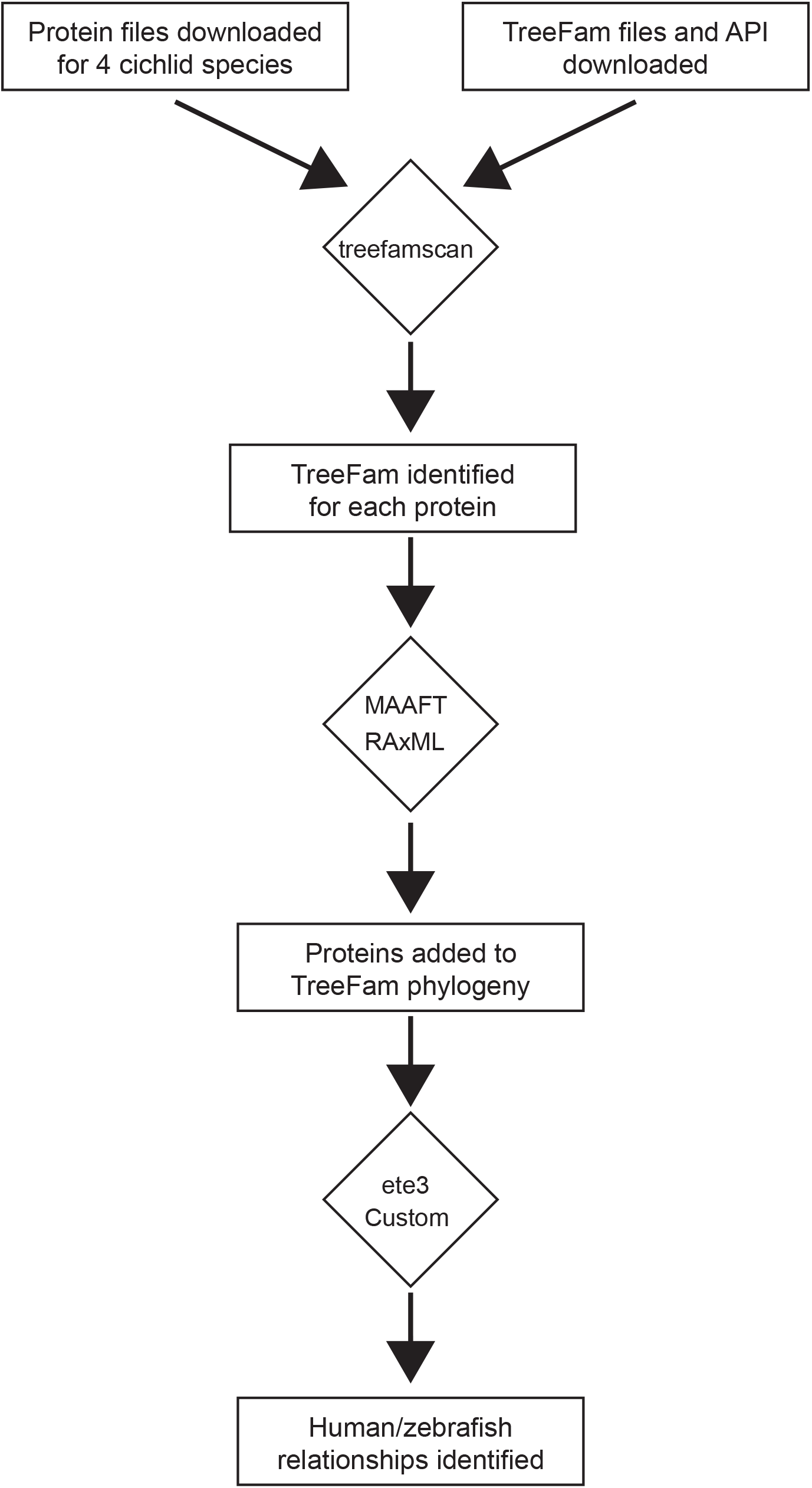
Pipeline for adding cichlid proteins to preexisting Treefam phylogenies

**Figure 2.**
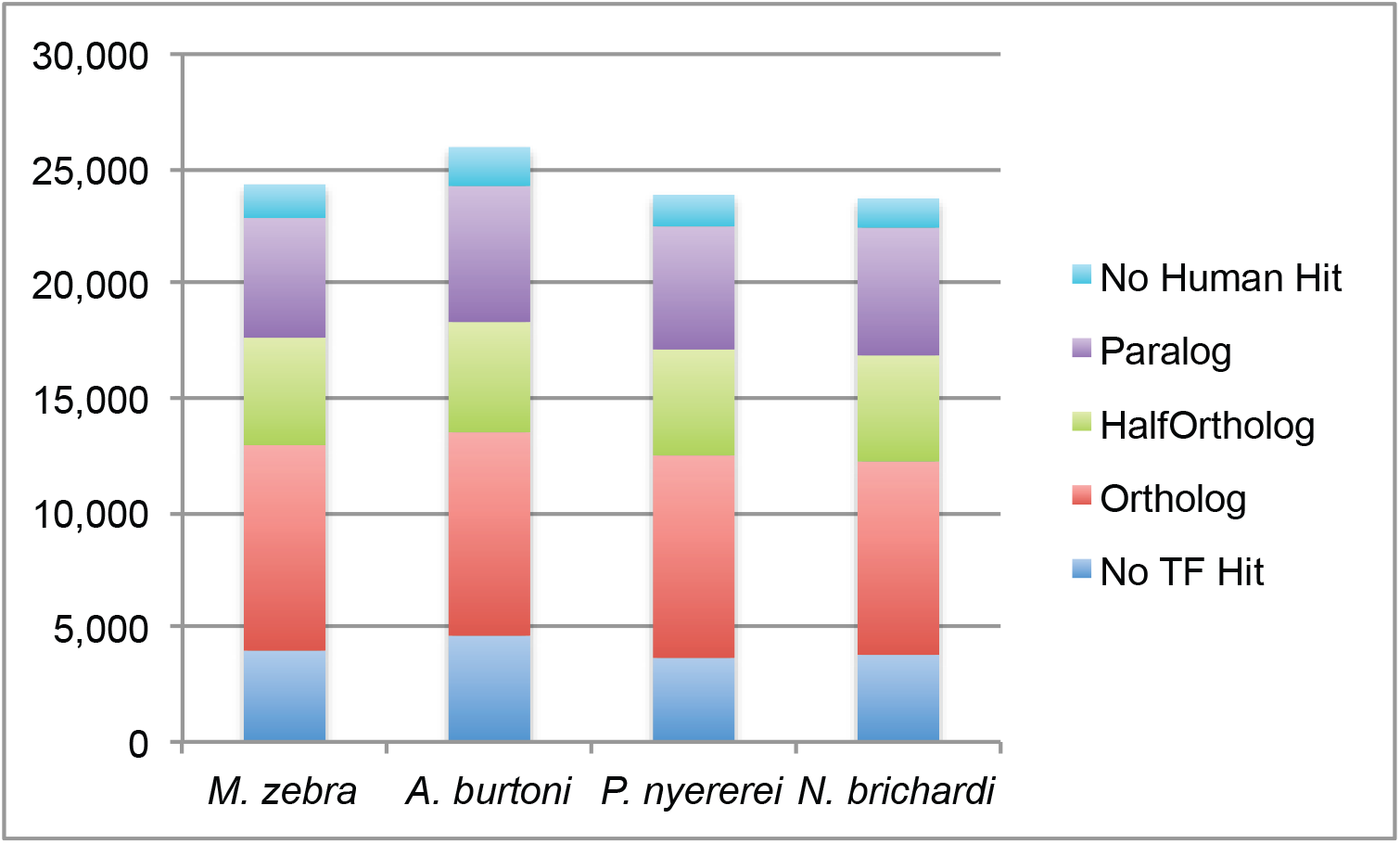
Summary of the human evolutionary relationships found in each species. HalfOrtholog is a non-standard relationship indicating a gene potentially duplicated and retained in an ancient teleost ancestor.

### Data accessibility

This data is intended as a resource for the cichlid community. We have provided access to this data in three ways. 1. Two PDF files for each TreeFam were generated for the purposes of human inspection. One PDF contains a phylogeny for a TreeFam from a limited number of species: humans, four well-characterized model organisms (*C. elegans, D. melanogaster, D. rerio,* and *M. musculus*), Nile tilapia (O. *niloticus),* and the four recently studied cichlid species. The second PDF contains a phylogeny of all 108 species used in the analysis. While the second phylogeny is the most complete, it is difficult to analyze due to the large number of species. This data is hosted on a web server (http://cichiids.biosci.gatech.edu/) and can be searched using cichlid gene names, TreeFam IDs, or human and zebrafish names. 2. Excel files for each cichlid species that contain each gene, its best hit to a human and zebra fish gene, and its evolutionary relationship to that gene. We anticipate this data will be useful for genomic scale analysis. For example, the excel file can be loaded into scripts to automatically map cichlid genes to human or zebrafish homologs. This could be useful for the purposes of pathway analysis (such as gene ontology), which often are limited to human genes. 3. Finally, phylogenies of each TreeFam are available for download in enhanced Newick tree format. These will be useful for any researchers interested in automated analysis of the phylogenies for the purpose of enhancing the evolutionary relationships that we have reported here. For example, researchers could use this dataset to identify genes whose protein phylogenies contradict the species phylogenies.

### Example phylogeny generated from a tree containing members of the TGFβ superfamily

To illustrate these evolutionary relationships as well as common issues users should be aware of in using these trees, we have included two figures of new phylogenies generated in this analysis. **Figure 3** shows a subtree of TF351789, which includes members of the TGFβ-superfamily of proteins including BMP2 and BMP4. These proteins are ubiquitous throughout metazoans, and control proliferation and differentiation of cells throughout development (Salazar, Gamer et al. 2016). This tree includes both ortholog and paralog relationships. For example, the subtree indicated by **a** in **Figure 3** shows ortholog relationships between the cichlid proteins and human BMP4. These genes likely play similar biological roles in cichlids. Similarly, subtree **b** contains cichlid orthologs to human BMP2 (with the exception of *M. zebra,* which will be discussed below), suggesting these genes play similar biological roles as the orthologs play in other species. There is also a cichlid-specific set of paralogs to BMP2 and BMP4 not present in *D. rerio* (subtree **c**) suggesting that there was a duplication of BMP2 or BMP4 in a recent common ancestor of all cichlid species following separation from the zebrafish lineage. It is not obvious from the phylogeny what biological role these genes might play. This clade of genes is potentially of interest to cichlid biologists, as they could play a role in the extensive morphological diversity observed among cichlid species. However, analysis of the full tree indicated that this clade contains genes from a large number of additional teleost fish along with a coelacanth fish (*L. chalumnae*) and an anole lizard (*A. carolinensis*) (**Figure S1**). Further, blasting the protein sequence encoded by the ab.gene.s112.4 from *A. burtoni* to the *D. rerio* genome identified a match to a known protein annotated as BMP16 (Feiner, Begemann et al. 2009). BMP16 does not appear to be present in the Treefam database, which explains why it was not present in the phylogeny. This set of BMP2/BMP4 paralogs thus seems to be a duplication that occurred in an ancient vertebrate ancestor of these fish (preceding the teleost ancestor) and lost in most tetrapod lineages as proposed by Marques et al (Marques, Fernandez et al. 2016).

**Figure 3.**
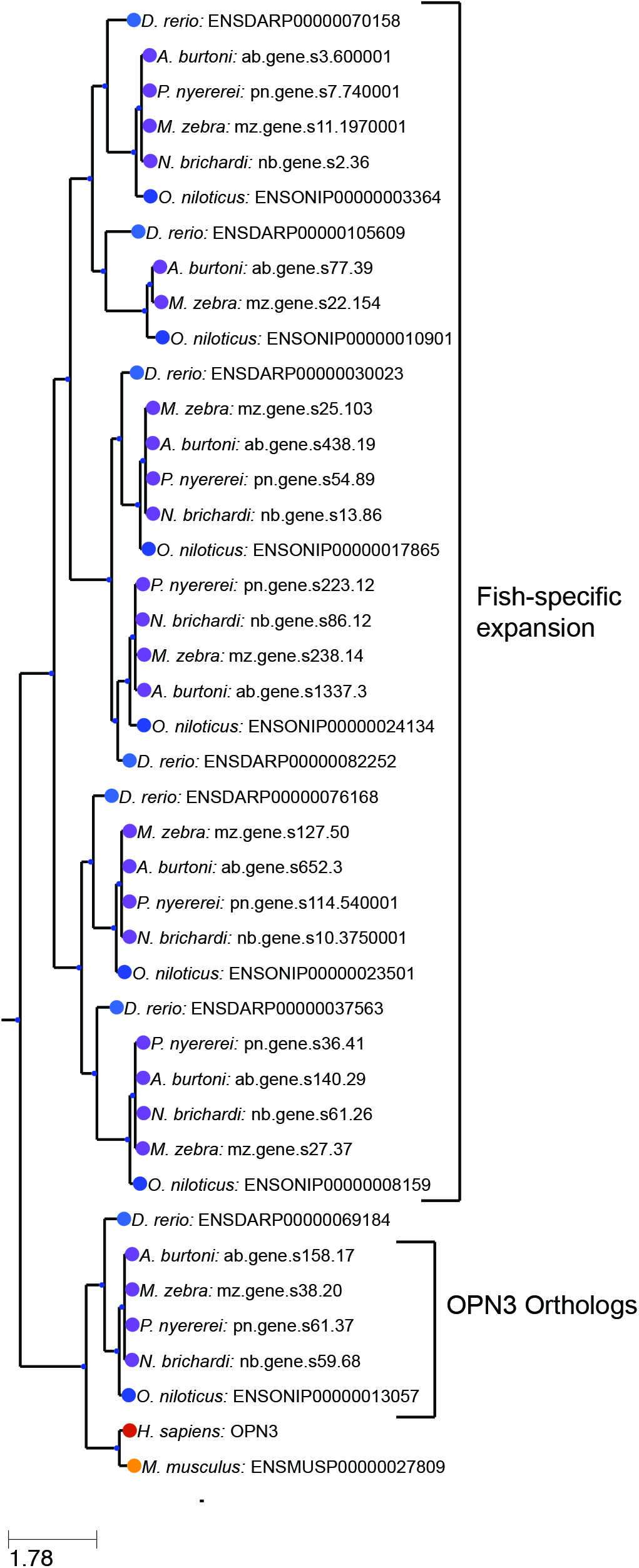
Subtree from the TF351789 family from a limited number of cichlid species and well-studied model organisms. This family contains a number of BMP growth factors belonging to the transforming growth factor beta family. Letters indicate additional subtrees discussed in the text.

We observed a similar issue in the phylogeny surrounding the human IRX1 gene (TF319371) (**Figure S2**). The Treefam phylogeny suggests that *D. rerio* contained a single ortholog to this gene while each of the cichlid species contained two copies of this gene. However, previous publications demonstrate that there are also two versions of IRX1 in *D. rerio* (called *irx1a* and *irx1b*) (Dildrop and Ruther 2004,Feijoo, Manzanares et al. 2004). Inspection of the Treefam data indicates that *irx1a* isn’t present in the starting dataset. These examples illustrate a common issue to most genomic analysis. Since Treefam relies on genomic-scale predictions, there are likely errors within the resulting phylogenies. Users would do well to manually verify or repeat any of these phylogenies for genes they are especially interested in.

We also were curious about the lack of a clear ortholog to BMP2 in *M. zebra* (**Figure 3**). It seemed unlikely that this species could lose this protein entirely due to its essential function in bone development. We were able to track down this discrepancy to an error in the annotation file for *M. zebra.* Through blastp, we were able to identify mz.gene.s5.238 as a gene containing a strong match to BMP2. mz.gene.s5.238, however, was assigned to the TF314677 family, and predicted to be an ortholog to the human protein FERMT1. When we invested the protein sequence more closely, it became clear that mz.gene.s5.238 appeared to contain a fusion of two genes: an ortholog to BMP2 and an ortholog to FERMT1. Due to the longer length of FERMT1, mz.gene.s5.238 was assigned to the TreeFam containing FERMT1. This is unlikely to represent a real gene fusion, and the improved version of the *M. zebra* genome predicts separate gene products consistent with other species (Conte and Kocher 2015). We observed a similar potential error with the PTGFR prostaglandin receptor. An ortholog of PTGFR has recently been shown to control female reproductive behaviors in the cichlid A. burton (Juntti, Hilliard et al. 2016)i, however, the Treefam containing the human PTGFR gene (TF324982), did not contain an ortholog of this gene in *A. burtoni.* Again, this seems to be due to an error in annotation incorrectly predicting a fusion between two genes. The best blastp match ab.gene.s495.12 contains a fusion between two genes, an ortholog to PTGFR and an ortholog to the ZFYVE9. Due to the longer length of the ZFYVE9 protein, the ab.gene.s495.12 gene is assigned to the Treefam containing the human ZFYVE9. Again, this is unlikely to represent a real fusion, and it since has been corrected in new annotations. These two examples illustrate how errors in the gene annotation can lead to incorrect phylogenies.

### Example phylogeny generated from a tree containing arginine vasopressin receptors

**Figure 4** shows a subtree of the phylogeny for TF106499, which contains a number of receptors for the arginine vasopressin and oxytocin neuropeptides that are thought to play a role in social behavior and sexual motivation (Hammock, Lim et al. 2005,Insel 2010). We have limited this phylogeny to the clade containing the AVPR1A and AVPR1B human proteins. The clade indicated by **a** demonstrates the HalfOrtholog relationship (**Figure 4**). All of the sequenced cichlid species (along with zebrafish and other teleost fish) contain two genes that fall within this clade. This phylogeny suggests that the function of the ancestral AVPR1A gene bifurcated into two genes in an ancestor to the teleost lineage. While the phylogeny suggests that both of these receptors should retain a molecular role in arginine vasopressin/oxytocin signaling, the biological function of AVPR1A should not be assigned to either of the two genes in each cichlid species. Rather, experiment will be necessary to parse out the biological function of each of these two half orthologs. A recent paper characterizing the expression pattern of these two receptors in zebrafish demonstrated that these two genes are expressed in similar but nonoverlapping cell types(Iwasaki, Taguchi et al. 2013). This phylogeny also contains the human AVPR1B protein. While mouse contains a clear ortholog to this gene, none of the cichlid species nor zebrafish contain an ortholog to this gene. Analysis of the full phylogeny suggests that AVPR1B was lost in the teleost fish completely. Thus, the phylogeny indicates that the biological functions assigned to AVPR1B through the study of mouse and other mammals should not be directly assigned to any of the cichlid homologs without experimental study.

**Figure 4.**
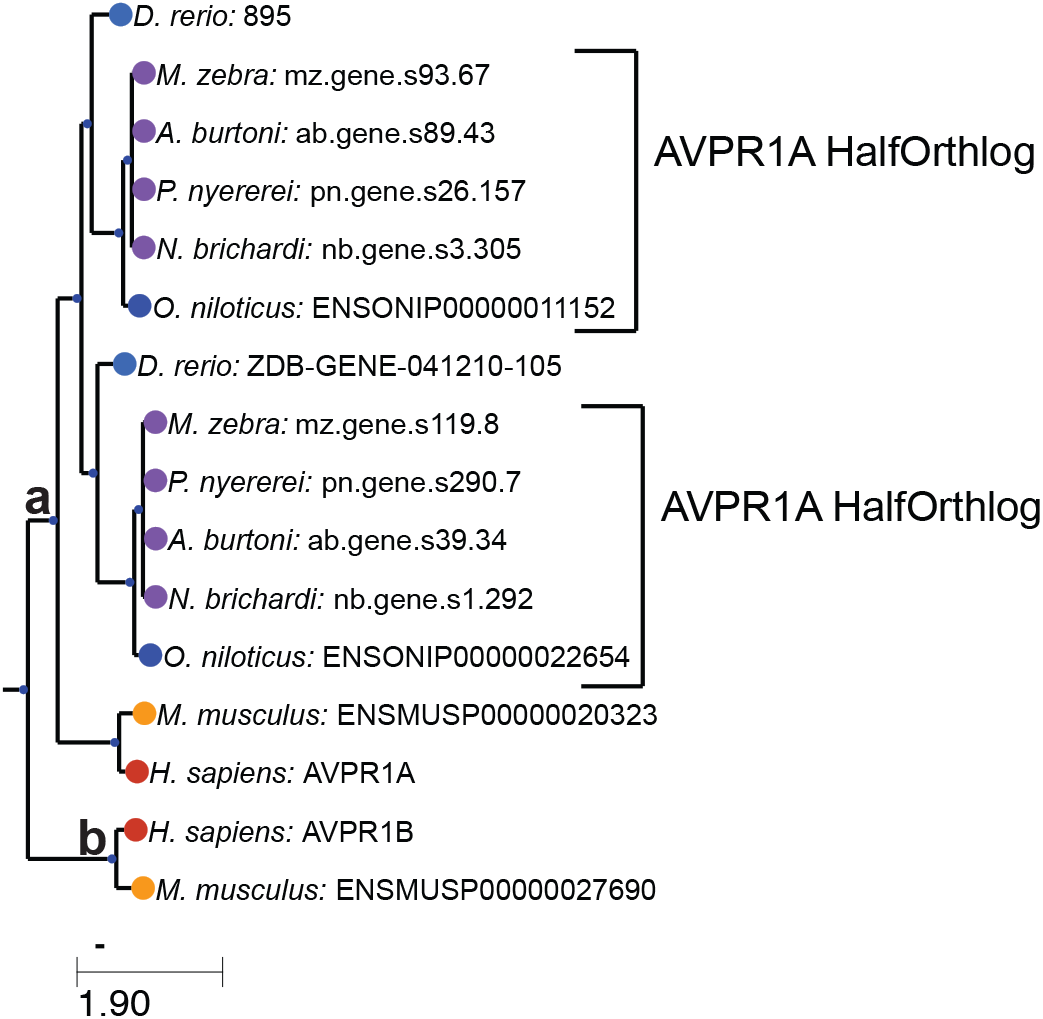
Subtree from the TF106499 family from a limited number of cichlid species and well-studied model organisms. This family contains a number of G-protein receptors for the arginine vasopressin and oxytocin nonapeptide hormones. Letters indicate additional subtrees discussed in the text.

## CONCLUSION

This study reports a set of protein phylogenies generated for four recently sequenced African cichlids. We hope that these phylogenies will be useful for cichlid researchers for the purpose of inferring biological and molecular function of cichlid genes.

## ACKNOWLEDGEMENTS

We thank members of the McGrath and Streelman labs for comments on this work. This work was supported by NIH grants R21AG050304, R01GM114170, and the Ellison Medical Foundation (to P.T.M.) and NIH grants 1R01GM101095 and 2R01DE019637 (to J.T.S.).

---

**Figure S1.** Full tree for the TF351789 family from all 109 species included in the Treefam database. This family contains a number of BMP growth factors belonging to the transforming growth factor beta family.

**Figure S2.** Subtree from the TF319371 family from a limited number of cichlid species and well-studied model organisms. This family contains a number of Iroquois-family of homeodomain transcription factors involved in patterning and other development processes.

